# Attenuated incubation of ethanol-induced conditioned taste aversion in a model of dependence

**DOI:** 10.1101/2023.09.13.557582

**Authors:** Lindsey A Ramirez, Kathryn R Przybysz, Joseph R Pitock, E Margaret Starr, Hyerim Yang, Elizabeth J Glover

## Abstract

**Rationale:** Preclinical studies report attenuated ethanol-induced conditioned taste aversion (CTA) following chronic ethanol exposure, suggesting that tolerance develops to the aversive properties of ethanol. However, these studies are confounded by pre-exposure to the unconditioned stimulus (US; ethanol), which is well known to hinder conditioning.

**Objectives:** This study was designed to determine whether chronic ethanol exposure produces tolerance to the aversive properties of ethanol in the absence of a US pre-exposure confound.

**Methods:** CTA was performed in adult male and female Long-Evans rats by pairing 0.1% ingested saccharin with an intraperitoneal injection of ethanol (1.5 or 2.0 g/kg) or saline. Rats were then rendered ethanol dependent using chronic intermittent ethanol (CIE) vapor exposure. Controls were exposed to room air (AIR). The effect of chronic ethanol on CTA expression and reconditioning were examined following vapor exposure.

**Results:** Prior to vapor exposure, both sexes developed CTA to a comparable degree with 2.0 g/kg producing greater CTA than 1.5 g/kg ethanol. Following vapor exposure, AIR controls exhibited an increase in CTA magnitude compared to pre-vapor levels. This effect was absent in CIE-exposed rats. These group differences were eliminated upon re-conditioning after vapor exposure.

**Conclusions:** These data suggest that chronic ethanol does not facilitate tolerance to the aversive properties of ethanol but rather, attenuates incubation of ethanol-induced CTA. Loss of CTA incubation suggests that CIE exposure disrupts circuits encoding aversion.

## Introduction

Though the majority of adults are able to consume alcohol in a controlled manner, approximately 10.6% of people above the age of 12 develop alcohol use disorder (AUD) (U.S. Department of Health and Human Services, Substance Abuse and Mental Health Services Administration, Center for Behavioral Health Statistics and Quality, 2021),characterized by heavy drinking that is often uncontrolled and persists despite significant negative consequences to health and well-being (American Psychiatric Association, 2013). Sensitivity to the rewarding (e.g., stimulation, euphoria) and aversive (e.g., sedation, nausea) properties of ethanol plays an important role in regulating continued drinking (Ray et al., 2010; Verendeev & Riley, 2013). For example, heavy drinkers consistently report experiencing higher levels of stimulation and lower levels of sedation than light drinkers (L. Holdstock et al., 2000; King et al., 2002, 2011, 2016; Morean & Corbin, 2010; Roche et al., 2014). In addition, a greater response to ethanol’s rewarding properties combined with a lower response to its aversive properties predicts AUD diagnosis (Haass-Koffler & Perciballi, 2020; Schuckit et al., 1988).

Pavlovian conditioning paradigms have been used to interrogate the subjective response to drugs in rodents, with conditioned place preference (CPP) most frequently used to measure a drug’s rewarding properties and conditioned taste aversion (CTA) used to measure its aversive properties. These paradigms use a novel context (CPP) or tastant (CTA) as a conditioned stimulus (CS) that is paired with drug administration (unconditioned stimulus; US). The amount of time spent in the drug-paired context or consuming the drug-paired tastant on test day is used as a measure of response to the drug’s rewarding or aversive properties, respectively. Using these approaches, preclinical studies have produced findings that parallel those observed in the clinic. For example, a significant negative relationship was observed between ethanol-induced CTA and preference for ethanol during home cage drinking in a study that characterized subjective response in 15 different mouse strains (Broadbent et al., 2002). A meta-analysis of both mouse and rat models of AUD reported similar results, with greater ethanol-induced CTA associated with lower levels of home cage ethanol consumption (Green & Grahame, 2008). Thus, findings from both human and rodent studies demonstrate the ability of ethanol’s rewarding properties to promote continued use and its aversive properties to limit use, similar to observations made of other addictive drugs (Davis & Riley, 2010; Verendeev & Riley, 2013).

Of note, evidence suggests that individuals chronically exposed to ethanol develop tolerance to its aversive properties (Corbin et al., 2013; Morean & Corbin, 2008), further releasing the brake on drinking. Findings from preclinical studies in which animals were repeatedly exposed to ethanol support these data. For example, alcohol-preferring P rats that drank ∼10 g/kg ethanol/day during continuous access home cage drinking exhibited attenuated ethanol-induced CTA compared to P rats that had access to water only prior to CTA testing (Stewart et al., 1991). Attenuated CTA has also been observed following use of well accepted models of dependence. This includes studies using mice or rats that received repeated ethanol administration via intragastric (i.g.) gavage (May et al., 2015), ethanol vapor (Diaz-Granados & Graham, 2007; Lopez et al., 2012), or intraperitoneal (i.p.) injection (Gauvin & Holloway, 1992). Interestingly, Cunningham et al. (2002) observed attenuated conditioned place aversion in the absence of any effect on CPP in mice that received repeated i.p. ethanol, suggesting that tolerance develops to the aversive, but not rewarding, properties of ethanol.

The strength of Pavlovian conditioning paradigms, such as CTA, depends on several factors. For example, exposure to either the US or CS prior to conditioning is well known to hinder learning the desired association (Chance, Paul, 2003; Hall & Symonds, 2006; Meyer et al., 2004; Revillo et al., 2013), including paradigms that use ethanol as the US. For example, a single pre-exposure to acute ethanol (i.g.) attenuated ethanol-induced CTA in male Sprague Dawley rats (Rabin et al., 1988). Similar findings were reported in male and female Long-Evans rats (Risinger & Cunningham, 1995; Sherrill et al., 2011) and male Holtzman albino rats (Barker & Johns, 1978) repeatedly exposed to ethanol prior to CTA. While tolerance could explain this effect for ethanol-induced CTA, attenuated learning is also observed in rats pre-exposed to other USs for which tolerance is not a concern (e.g., irradiation, LiCl) (Rabin et al., 1988) as well as in Pavlovian conditioning of non-drug associations (Björkstrand, 1990; Chang et al., 2007). Therefore, US pre-exposure effect is important to consider given that each of the studies showing tolerance to the aversive properties of ethanol in rodents measures ethanol-induced CTA after repeated pre-exposure to the US. Consequently, these studies interpret attenuated CTA as tolerance despite being indistinguishable from attenuated CTA observed following US pre-exposure.

By developing a new model that eliminates the US pre-exposure confound, the present study was designed to clarify whether attenuated ethanol-induced CTA following dependence is indicative of tolerance to the aversive properties of ethanol. Our results reveal an unexpected loss of CTA incubation in adult male and female Long-Evans rats chronically exposed to ethanol, suggesting that attenuated CTA previously reported in the literature results from ethanol pre-exposure and not tolerance per se. Our findings highlight the importance of eliminating US pre-exposure effects in future studies and the need for additional research using rigorous models to explore the effect of chronic ethanol exposure on behavioral responding to aversive stimuli.

## Materials & Methods

### Animals

All experiments were conducted in accordance with the National Institutes of Health Guide for the Care and use of Laboratory Animals and were approved by the University of Illinois Chicago Institutional Animal Care and Use Committee. Female and male adult Long-Evans rats (∼P60 on arrival; Envigo, Indianapolis, IN) were singly housed for the duration of the experiment in standard polycarbonate cages in a temperature and humidity-controlled room. Lights in the colony room were on a 12-hour reverse light/dark cycle (lights off at 10:00). Rats were allowed to acclimate for at least one week following arrival. Rats had *ad libitum* access to standard laboratory chow (Teklad 7912, Envigo) for the duration of the experiment and to water during the acclimation period. **Figure 1** depicts the overall experimental design described in detail below.

**Figure 1.**
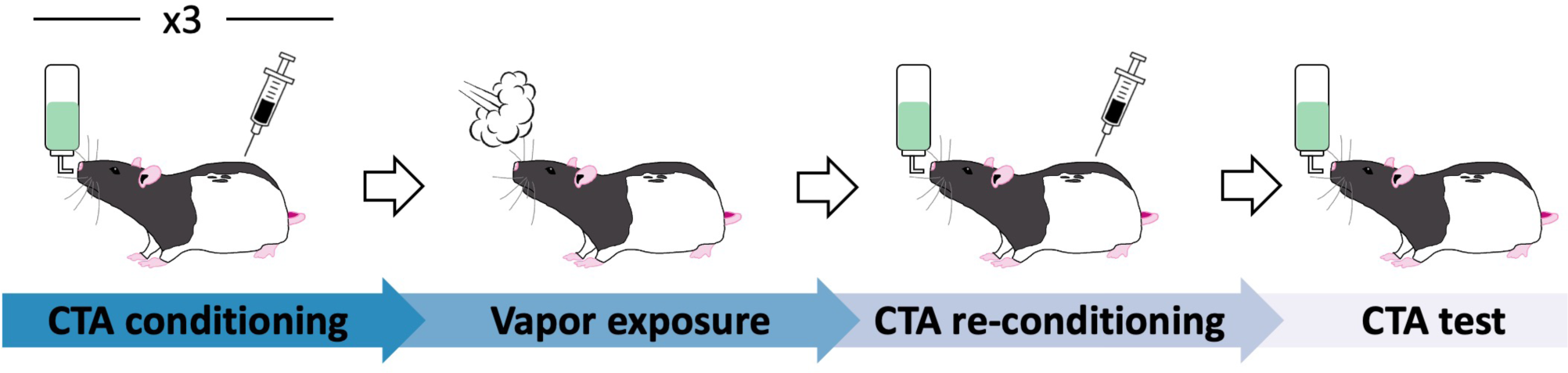
Improved experimental design eliminates the US pre-exposure confound. Rats first underwent a standard conditioned taste aversion (CTA) paradigm during which they were given access to 0.1% saccharin followed by i.p. injection of either EtOH (1.5 g/kg or 2.0 g/kg) or saline. After three conditioning trials, rats were rendered dependent using a standard chronic intermittent ethanol vapor exposure paradigm. A within-subjects design was used to evaluate change in CTA expression. Using this approach, rats were tested for CTA expression after vapor exposure and then re-conditioned by pairing saccharin with drug injection again. This was followed by the final saccharin access period during which saccharin was not paired with drug injection.

### Conditioned taste aversion

CTA was performed prior to chronic ethanol exposure as previously described (Glover et al., 2016; Przybysz et al., 2023). Briefly, rats were habituated to scheduled fluid access for one week during which they were given tap water for 30 minutes daily. Next, rats underwent three conditioning trials to associate a 0.1% saccharin solution (Sigma Aldrich, Missouri, in tap water) with drug injection. During each conditioning trial, rats were provided with saccharin during the 30-minute access period after which they received an injection (i.p.) of one of two doses of ethanol (EtOH; 1.5 g/kg or 2.0 g/kg; 20% v/v, Decon Laboratories, Pennsylvania in ddH_2_O) or saline (EtOH equivalent volume; Baxter Healthcare Corporation, Illinois). Rats were assigned to drug groups in a manner counterbalanced for baseline saccharin intake obtained on conditioning day 1. Each conditioning trial was followed by at least three water recovery days identical to scheduled access during the habituation period. Fluid intake was measured daily by calculating change in bottle weight. Spillage was controlled for during each experimental session by correcting change in bottle weight for average fluid loss from bottles placed on an empty cage.

### Chronic intermittent ethanol exposure

After the third CTA conditioning day, rats were rendered dependent using a standard 14-day chronic intermittent ethanol (CIE) vapor exposure paradigm using previously published procedures (Glover et al., 2019, 2021). Rats were assigned to receive exposure to either ethanol vapor (CIE) or room air (AIR) for 14 hours per day for 14 consecutive days in a manner counterbalanced by drug group (n=12-13/sex/group). Rats had *ad libitum* access to water during this time. Rats were scored for behavioral signs of intoxication immediately after each 14-hour vapor exposure session using a subjective rating scale with scores ranging from 1 (no signs of intoxication) to 5 (loss of consciousness) (Glover et al., 2019, 2021). Blood was collected via tail nick from rats exposed to ethanol vapor on four separate occasions during the 14-day exposure period. Because ethanol intoxication can produce analgesic effects, air-exposed controls received a tail pinch in lieu of a tail nick to mimic a similar level of temporary discomfort experienced by the ethanol-exposed rats during tail nick.

### CTA re-conditioning & test

Rats were left undisturbed for three days following their final vapor exposure session during which they had *ad libitum* access to water. To examine the effect of chronic ethanol exposure on ethanol-induced CTA expression, scheduled fluid access resumed on the fourth day after their final vapor exposure session. After three days of scheduled water access, rats were once again given saccharin followed by drug injection. Re-conditioning (conditioning day 4) was followed by at least three water recovery days. During the final test, rats were provided with saccharin for the standard 30 min access period but on this day no drug was administered.

### Blood ethanol concentration

Blood samples were centrifuged (10,000 rcf, 10 min, 4°C) and the plasma collected and stored at -20°C until ready for analysis. Blood ethanol concentration (BEC) was measured using an Analox AM1 Analyzer (Analox Instruments; Atlanta, GA).

### Statistical analysis

Two distinct phenotypes emerged during pre-vapor CTA testing – one phenotype that was relatively sensitive to ethanol-induced CTA and another that was relatively resistant. Differences between these two phenotypes in CTA expression prior to vapor exposure were reported in a previous manuscript (Przybysz et al., 2023). Because the analysis performed in (Przybysz et al., 2023) was limited to data prior to vapor exposure, it did not incorporate future vapor group assignment into the analysis as has been performed in the current manuscript. Thus, the pre-vapor CTA data reported in the present manuscript constitutes a new analysis of data previously reported in (Przybysz et al., 2023). Importantly, we observed no significant effect of phenotype on CTA after vapor exposure (**See supplemental material**). Therefore, phenotype is not included as a factor in the analyses that follow.

Saccharin intake (mL) was used as a measure of CTA with low intake indicating strong CTA. Outlier analysis was performed on baseline saccharin intake using the ROUT method, which revealed the presence of one outlier. Thus, intake on conditioning day 1 was excluded for one female in the AIR 2.0 g/kg EtOH group. Additionally, saccharin intake on conditioning day 3 was excluded for one female in the AIR saline group due to technical error. All other data points for these rats were included.

Normality tests were performed in GraphPad Prism. Data for saccharin intake and intoxication were normally distributed. Levene’s test was used to measure the homogeneity of variance for saccharin intake. The Geisser-Greenhouse correction was used when equality of variances was not assumed. Saccharin intake across the first three conditioning days (prior to vapor exposure) was analyzed using a multifactorial ANOVA. Between group differences in saccharin intake at baseline and comparisons of saccharin intake at distinct timepoints were analyzed using two-way ANOVA. BECs and intoxication scores were analyzed using one-way ANOVAs. Pearson correlations were performed to examine the relationship between BECs and change in saccharin intake prior to and following vapor exposure. Data were analyzed using GraphPad Prism (version 9.2.0) or SPSS (Version 29.0.0.0 [241]). Data are presented as mean ± SEM and were considered statistically significant at p ≤ 0.05.

## Results

### Ethanol-induced CTA is similar in male and female rats prior to vapor exposure

A multifactorial repeated measures ANOVA was used to determine the effect of sex, drug group, and vapor group on saccharin intake across conditioning days. This analysis revealed a significant main effect of sex [*F*(1,143= 58.14, p<0.001]. Bonferroni-corrected multiple comparisons revealed that males consumed significantly more saccharin than females (p<0.001). Therefore, data from each sex were analyzed separately in all subsequent analyses.

Rats were assigned to drug and vapor groups in a manner counterbalanced for baseline saccharin consumption. This was confirmed using two-way ANOVAs, which found no significant between-group differences in saccharin intake on conditioning day 1 in either males [Drug: *F*(2,72)=0.01, *p*=0.99; Vapor: *F*(1,72)=0.59, *p*=0.44; Interaction: *F*(2,72)=1.08, *p*=0.34; **Figure 2A**] or females [Drug: *F*(2,70)=0.32, p=0.73; Vapor: *F*(1,70)=0.05, p=0.82; Interaction: *F*(2,70)=0.10, p=0.91; Figure 2C].

**Figure 2.**
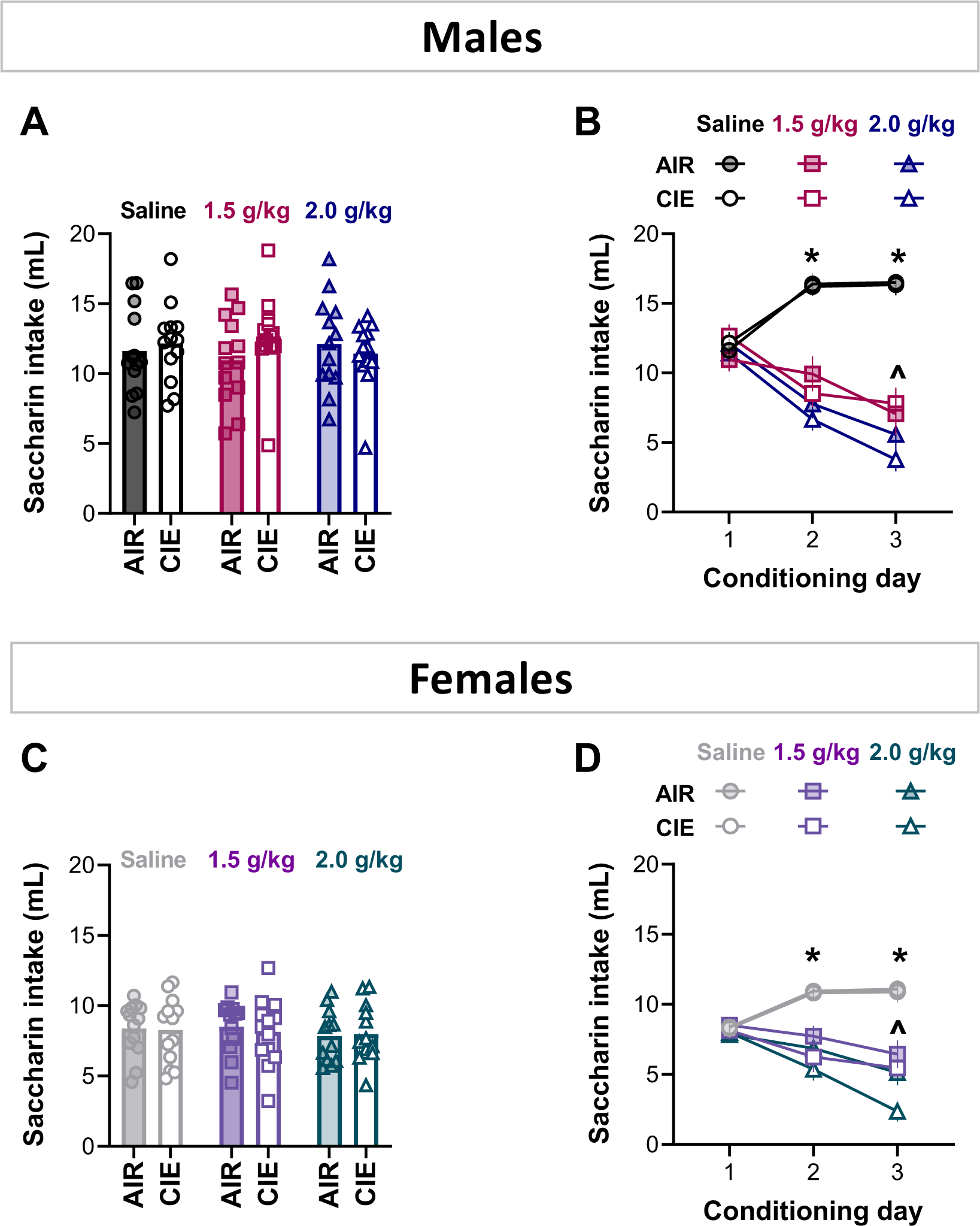
Ethanol produces dose-dependent CTA in both male and female rats. No significant differences in saccharin intake were observed at baseline (conditioning day 1) in male **(A)** or female **(C)** rats. Males **(B)** and females **(D)** developed significant ethanol-induced CTA that was dose-dependent with greater CTA evident in rats that received 2.0 compared to 1.5 g/kg EtOH. n=12-13 rats per group; *p<0.05 EtOH groups vs saline; ^p<0.05 1.5 g/kg vs 2.0 g/kg.

Next, a multifactorial repeated measures ANOVA was used to compare saccharin intake between groups across conditioning days in males. No significant three way interaction was observed between drug group, vapor group, and conditioning day [*F*(3.65, 131.38)=0.79, *p*=0.52]. However, this analysis found significant main effects of conditioning day [*F*(1.83,0.79)= 17.92, p<0.001] and drug group [*F*(2,72)=46.50, *p*<0.001], as well as a significant interaction between drug group and conditioning day [*F*(3.65,0.79)=44.72, *p*<0.001]. Bonferroni-corrected multiple comparisons revealed that rats in both EtOH groups drank significantly less saccharin on conditioning days 2 and 3 compared to saline (p<0.001). In addition, rats in the 2.0 g/kg EtOH group drank significantly less saccharin than rats in the 1.5g/kg EtOH group on conditioning day 3 (p=0.03; **Figure 2B**).

Similarly, there was no significant three-way interaction between drug group, vapor group, and conditioning day in female rats [*F*(3.65,129.50)=0.42, p=0.79]. There were, however, main effects of drug group [*F*(2,71)=23.51, p<0.001] and conditioning day [*F*(1.82,129.50)=9.64, p<0.001], as well as a significant interaction between drug group and conditioning day [*F*(3.65,129.50)=20.61, p<0.001]. Similar to our observations in males, Bonferroni-corrected multiple comparisons found that rats in the EtOH groups drank significantly less saccharin on conditioning days 2 and 3 compared to saline controls (p<0.001) and rats that received 2.0 g/kg EtOH drank significantly less saccharin than rats that received 1.5g/kg EtOH on conditioning day 3 (p=0.02; **Figure 2D**). Altogether, these results show that the magnitude of ethanol-induced CTA is dose-dependent and is similar in male and female Long-Evans rats.

### Chronic intermittent ethanol exposure attenuates incubation of ethanol-induced CTA

To determine the impact of CIE exposure on ethanol-induced CTA, we performed *a priori* analyses comparing CTA expression prior to vapor exposure (conditioning day 3) to CTA expression after vapor exposure (conditioning day 4) within each drug group.

In male rats injected with saline, a two-way RM ANOVA found a significant interaction between vapor group and time [*F*(1,24)=32.28, *p*<0.0001, **Figure 3A**]. Sidak’s pairwise comparisons revealed that saccharin intake increased after vapor exposure in the CIE-exposed (*p*<0.0001), but not AIR-exposed (p=0.11), rats. A significant main effect of time on saccharin intake was observed in male rats injected with 1.5 g/kg EtOH, with saccharin intake decreasing following vapor exposure [*F*(1,24)=5.23, *p*=0.03, **Figure 3B**]. A significant interaction between vapor group and time was also observed in rats injected with 2.0 g/kg EtOH [*F*(1,24)=4.56, *p*=0.04, **Figure 3C**]. Sidak’s pairwise comparisons of this interaction revealed that AIR-exposed rats exhibited a significant decrease in saccharin intake after vapor exposure (p=0.006) that was absent in the CIE-exposed group (p=0.96).

**Figure 3.**
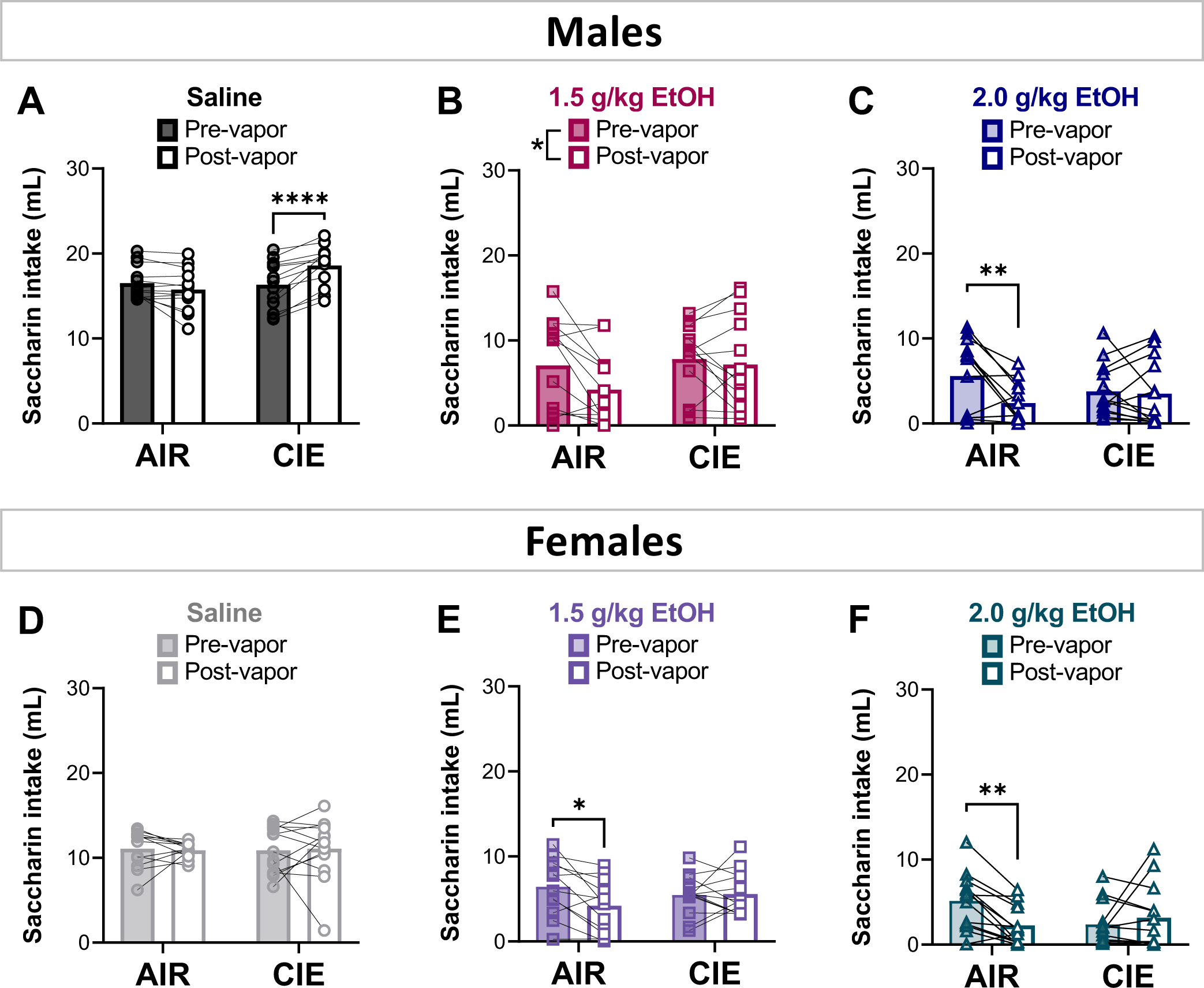
CIE exposure attenuates incubation of ethanol-induced CTA in male and female rats. In rats that received saline injections, saccharin intake significantly increased after vapor exposure in males in the CIE-exposed group **(A)**, whereas saccharin intake remained unchanged in females **(D)**. CTA increased in strength in male **(B-C)** and female **(E-F)** AIR-exposed controls that received either dose of EtOH – an effect that was absent in CIE-exposed counterparts. n=12-13 rats per group; *p<0.05; **p<0.01, ****p<0.0001.

The same analyses in females produced comparable results. No significant differences in saccharin intake were observed in saline-injected female rats when comparing saccharin intake before and after vapor exposure [Vapor: *F*(1,24)=8.40e-007, p=1.00; Time: *F*(1,24)=0.000, p>0.99; Interaction: *F*(1,24)=0.13, p=0.72; **Figure 3D**]. A significant interaction between vapor exposure and time was observed in female rats that had saccharin paired with 1.5 g/kg EtOH [F(1,23)=4.94, p=0.04]. Sidak’s multiple comparisons found that saccharin intake decreased significantly following AIR (p=0.02), but not CIE (p=0.98) exposure (**Figure 3E**). Similarly, a significant interaction between vapor exposure and time was observed in rats injected with 2.0 g/kg EtOH [*F*(1,24)=10.71, p=0.003]. Sidak’s multiple comparisons of this effect showed that, similar to findings in females that received 1.5 g/kg EtOH, saccharin intake was significantly lower after vapor exposure in the AIR-(p=0.003) but not CIE-exposed (p=0.54) group (**Figure 3F**).

### Levels of intoxication during vapor exposure are similar across drug groups

Behavioral signs of intoxication and blood ethanol concentrations (BECs) were similar across drug groups during vapor exposure. This was confirmed in male rats using a two-way RM ANOVA, which uncovered a significant main effect of time [*F*(7.31,261.4)=3.15, p=0.003] in the absence of any other significant effects. Dunnett’s post hoc analysis found that average intoxication scores were significantly higher on the first day of vapor exposure compared to days two and three (p<0.05; **Figure 4A**). Visual inspection of these data suggests that this effect is driven by high behavioral signs of intoxication in the saline group on day one compared to rats in the EtOH groups. One-way ANOVAs comparing average level of intoxication score and BEC in males across the 14 day vapor exposure period found no significant between-group differences in either measure [intoxication score: *F*(2,36)=0.44, *p*=0.65, **Figure 4B**; BEC:*F*(2,36)=1.85, *p*=0.17, **Figure 4C**].

**Figure 4.**
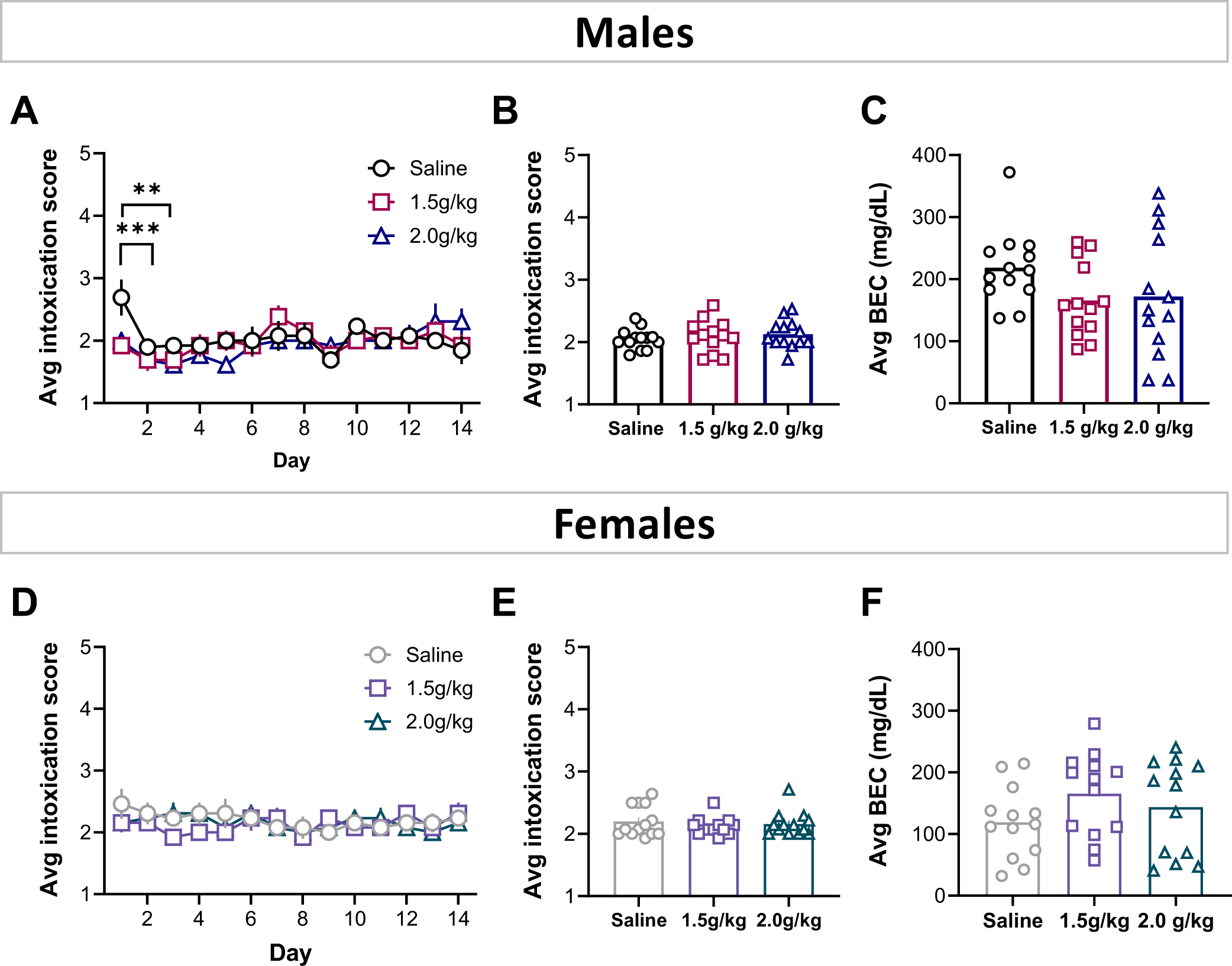
Intoxication levels did not differ between drug groups. Intoxication scores measured after daily ethanol vapor exposure differed significantly in males on day one compared to days two and three of the exposure period **(A).** This effect appears to be driven by unusually high behavioral signs of intoxication in rats that had saccharin paired with saline. In males, no other significant between-group differences were observed in intoxication scores **(B)** or BECs **(C)** averaged across all vapor exposure days. No significant between-group differences were observed in daily intoxication scores **(D)**, average intoxication scores **(E)** or average BECs **(F)** in females. n=12-13 rats per group; Asterisks indicate significant post hoc comparisons of the main effect of time; **p<0.01, ***p<0.001.

Similar results were found in females. A two-way RM ANOVA found no significant between-group differences in intoxication scores across vapor exposure days (all p values >0.30; **Figure 4D**). In addition, one-way ANOVAs revealed no significant differences in average intoxication score [*F*(2,36)=0.50, p=0.61, **Figure 4E**] or BEC [*F*(2,36)=1.53, p=0.23; **Figure 4F**] in female rats during the 14 day exposure period.

### Post-vapor reconditioning eliminates differences in CTA magnitude between AIR- and CIE- exposed rats

To determine whether ethanol-induced CTA was permanently attenuated in CIE-exposed rats, we performed *a priori* comparisons between CTA magnitude on conditioning day 4 (following vapor exposure, but prior to re-conditioning) with CTA magnitude after reconditioning within each drug group. In male rats that were injected with saline, a two-way RM ANOVA uncovered significant main effects of vapor group [*F*(1,24)=7.3, p=0.01] and time [*F*(1,24)=5.00, *p*=0.03] in the absence of any significant interaction between these variables [*F*(1,24) =0.41, p=0.53, **Figure 5A**]. These results show that CIE-exposed rats exhibited significantly greater saccharin intake than AIR-exposed controls regardless of conditioning timepoint. Additionally, saccharin intake increased significantly from conditioning day 4 to test day in both AIR- and CIE-exposed rats. In male rats injected with 1.5g/kg EtOH, saccharin intake was not significantly different between groups, although there was a trend for a main effect of time [*F*(1,24)=3.27, p=0.08; **Figure 5B**). A significant main effect of time [*F*(1,24)=15.17, p=0.0007] was observed in the absence of any other significant effects in males that received saccharin paired with 2.0 g/kg EtOH (**Figure 5C**) revealing that saccharin intake decreased from conditioning day 4 to test in both groups.

**Figure 5.**
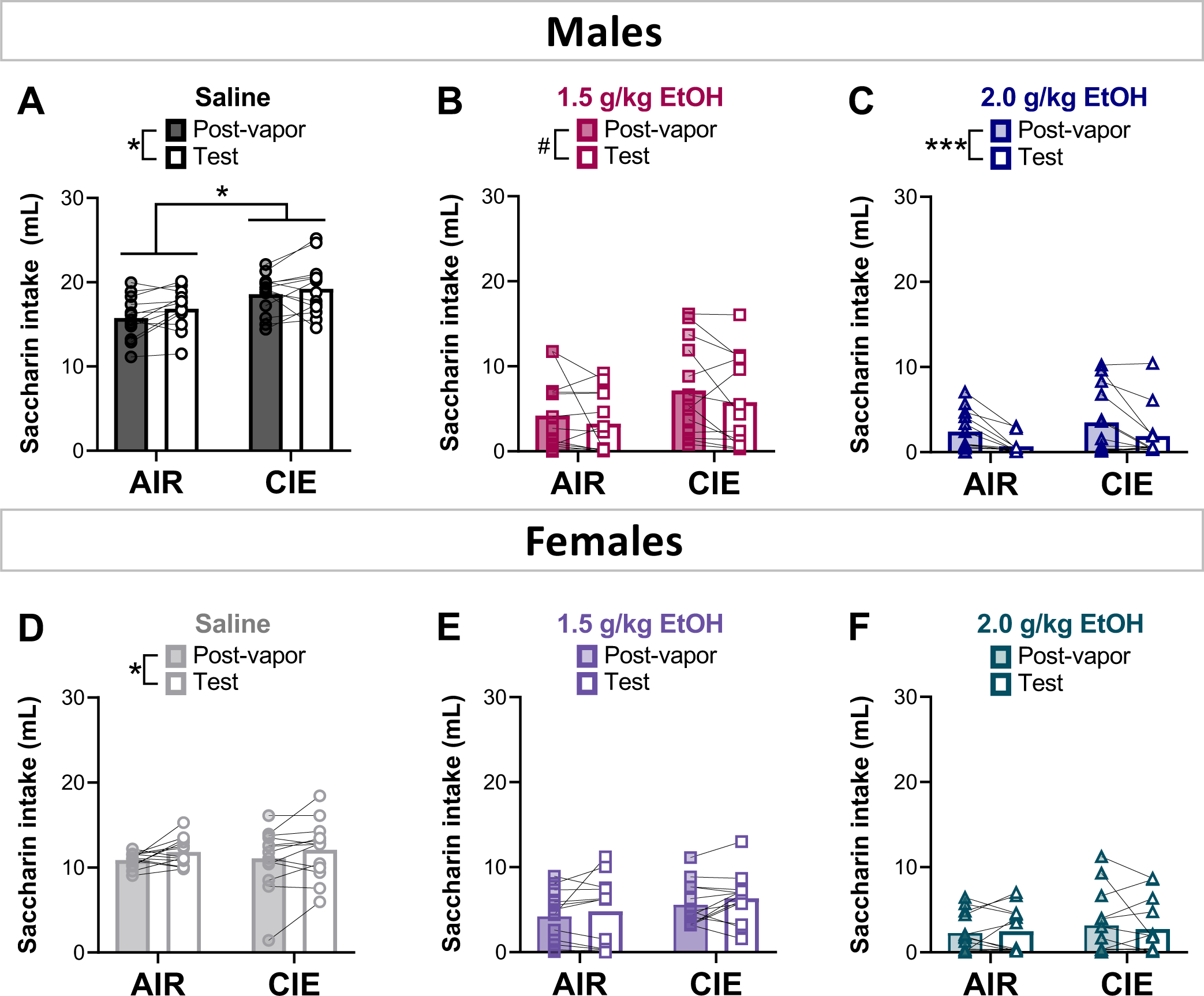
Re-conditioning eliminates post-vapor differences in CTA magnitude. Saccharin intake increased after re-conditioning in both male **(A)** and female **(D)** rats that had saccharin paired with a saline injection. Re-conditioning increased the strength of ethanol-induced CTA to a similar degree in both AIR- and CIE-exposed males. This effect trended toward significance in males that were injected with 1.5 g/kg EtOH **(B)** but reached statistical significance in males injected with 2.0 g/kg EtOH **(C)**. No between-group differences in CTA magnitude were observed in females injected with either 1.5 g/kg **(E)** or 2.0 g/kg **(F)** EtOH. n=12-13 rats per group; ^#^p=0.08; *p<0.05, ***p<0.001.

The same analysis in female rats injected with saline revealed a significant main effect of time [*F*(1,24)=7.59, p=0.01] with saccharin intake increasing significantly in both AIR- and CIE-exposed rats from conditioning day 4 to test day (**Figure 5D**). No significant differences in saccharin intake were observed between conditioning day 4 and test in rats injected with either 1.5 g/kg EtOH (all p values > 0.10; **Figure 5E**) or 2.0 g/kg EtOH (all p values > 0.50; **Figure 5F**).

### Relationship between intoxication during vapor exposure and post-vapor CTA magnitude

To explore whether the degree of intoxication during vapor exposure was related to change in CTA magnitude, Pearson correlations were used to analyze the relationship between average BEC during CIE exposure and percent change in saccharin intake from conditioning day 3 to 4. While a positive relationship was observed between these two measures in males that had saccharin paired with saline (r=0.38, p=0.21) or 1.5 g/kg EtOH (r=0.47, p=0.11), these correlations did not reach statistical significance (**Figure 6A-B**). This contrasts with males that had saccharin paired with 2.0 g/kg EtOH, which exhibited a statistically significant positive relationship between average BEC and change in saccharin intake (r=0.66, p=0.01; **Figure 6C**). Similar to males of the same drug groups, a nonsignificant positive relationship was observed between average BEC and change in saccharin intake in females that had saccharin paired with saline (r=0.14, p=0.64, **Figure 6D**) or 1.5 g/kg EtOH (r=0.40, p=0.18, **Figure 6E**). Unlike males, however, these variables were negatively correlated in females that received saccharin paired with 2.0 g/kg EtOH, although this relationship did not reach statistical significance (r=-0.25, p=0.42, **Figure 6F**).

**Figure 6.**
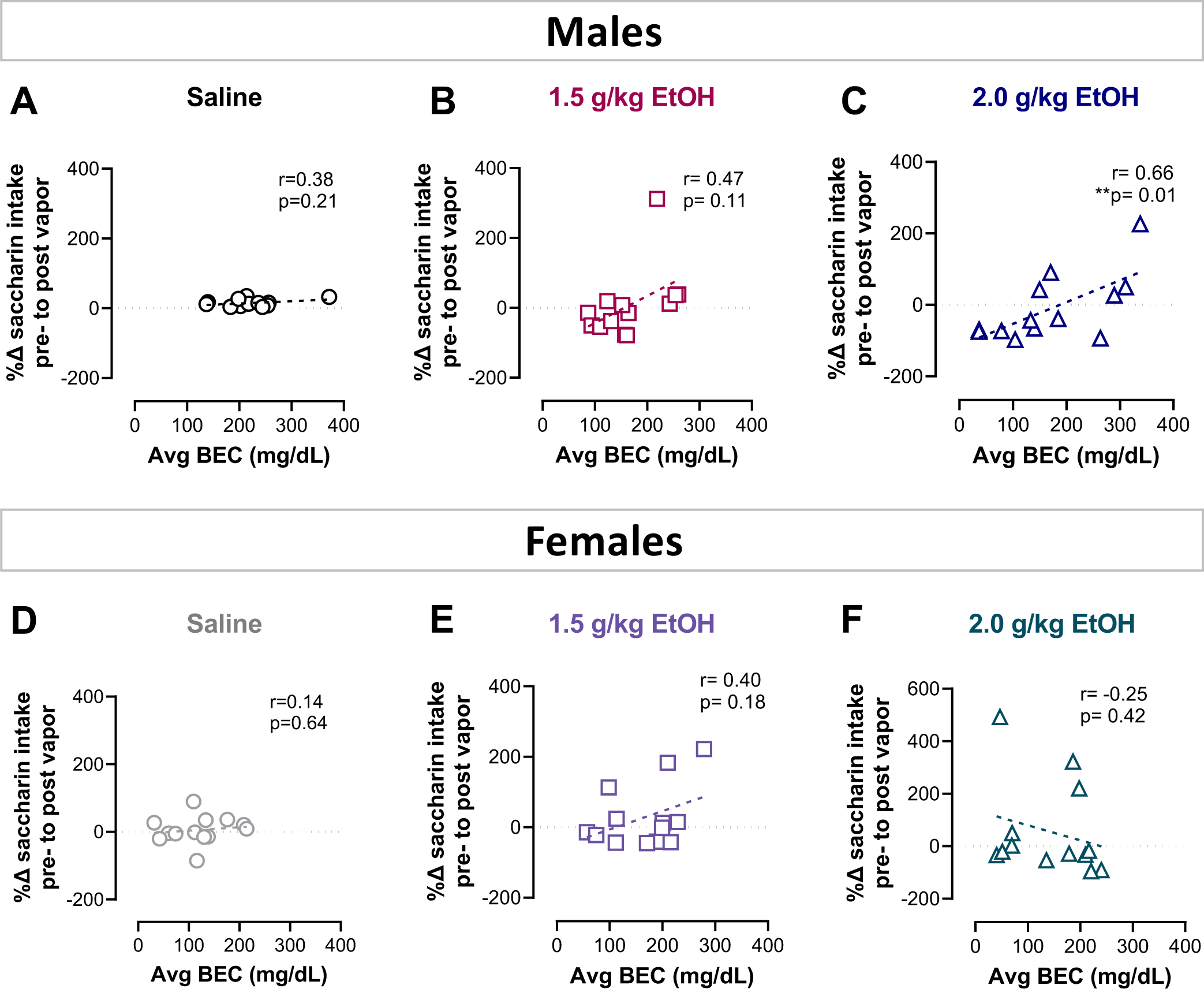
Relationship between intoxication and CTA magnitude. Pearson correlations were performed between average BEC and the percent change in saccharin intake prior to and following CIE exposure. There was no significant relationship between these variables in males injected with saline **(A)** or 1.5 g/kg EtOH **(B)**. Males injected with 2.0 g/kg EtOH **(C)** showed a statistically significant positive correlation between BEC and change in saccharin intake. Females that were injected with saline **(D)**, 1.5 g/kg EtOH **(E)**, and 2.0 g/kg EtOH **(F)** showed no correlation between these variables. n=12-13 rats per group; **p≤0.01.

## Discussion

Previous studies using CTA to examine tolerance to the aversive properties of ethanol are limited to between-subject comparisons of controls and experimental animals and are therefore confounded by the US pre-exposure effect. In contrast, the present study improves upon prior work by employing a within-subjects design to examine ethanol-induced CTA before and after chronic ethanol exposure, thereby eliminating the US pre-exposure confound. Using this approach, our results revealed a strengthening of ethanol-induced CTA in AIR-exposed male and female controls that is absent in CIE-exposed rats. These between-group differences in CTA did not persist following re-conditioning after vapor exposure. These findings suggest that CIE exposure does not produce tolerance to the aversive properties of ethanol, but rather, attenuates incubation of ethanol-induced CTA.

Incubation of conditioned fear, defined as an increase in the strength of conditioned responding to cues predictive of threatening stimuli (e.g., shock), is an often overlooked, but relatively well-characterized, phenomenon (Eysenck, 1968; McAllister & McAllister, 1967; Pickens et al., 2010). Previous studies in rodents have found that incubation of conditioned fear begins to emerge as early as 15 days after conditioning (Pickens, Golden, et al., 2009), is particularly robust 30 days after conditioning (Pickens, Adams-Deutsch, et al., 2009; Pickens et al., 2010; Pickens, Golden, et al., 2009; Pickens & Theberge, 2014), and can persist until at least 60 days after conditioning (Pickens, Golden, et al., 2009). Moreover, incubation occurs regardless of whether the aversive stimulus is avoidable or unavoidable (McMichael, 1966). Similar results have also been observed in humans (Bindra & Cameron, 1953; Sandin & Chorot, 1989). This phenomenon is likely an adaptive response to aversive conditioning that strengthens learning to facilitate prolonged avoidance of potentially harmful environmental stimuli.

While fear conditioning is much more broadly used by researchers than CTA, many parallels exist between the two Pavlovian paradigms. This includes data showing incubation of CTA similar to what is observed in conditioned fear. These results show moderate LiCl-induced CTA 1-2 days after conditioning that weakens within the first week after conditioning, followed by an increase in strength thereafter (Biederman et al., 1974; Marcant et al., 1985). Notably, both studies observed strong incubation of CTA 14 days after conditioning, which corresponds to the duration of vapor exposure in the present study. Thus, data from AIR-exposed controls replicate this previous work by showing strengthened CTA ∼ 14 days after the last conditioning trial. Our data also expands upon these findings by showing that incubation occurs for ethanol-induced as well as LiCl-induced CTA. Of note, CIE-exposed rats exhibited similar BECs and behavioral signs of intoxication throughout the vapor exposure period. In addition, no consistent relationship was observed between BEC and saccharin intake following vapor exposure. Therefore, it is unlikely that differences in level of intoxication contributed to differences in CTA magnitude after CIE exposure.

Chronic ethanol exposure is associated with a number of cognitive impairments including deficits in memory recall (Antonucci et al., 2020; Beracochea & Jaffard, 1985; Bernardin et al., 2014; Sachdeva et al., 2016; Walker & Hunter, 1978). With this in mind, we included a re-conditioning trial after vapor exposure to allow for consideration of whether the attenuated CTA we expected to see in CIE-exposed rats was due to impaired recall of the saccharin-ethanol association. Importantly, we did not observe attenuated recall of the learned association as saccharin intake prior to and following vapor exposure were not significantly different in CIE-exposed rats. Nevertheless, re-conditioning after vapor exposure eliminated differences in CTA magnitude between AIR- and CIE-exposed groups suggesting that a single reminder trial can restore CTA magnitude to control levels in CIE-exposed rats. Altogether, these data suggest that CIE exposure produces a transient loss of CTA incubation observed in AIR controls rather than tolerance to the aversive properties of ethanol that has been previously described.

In the present study, we found that CTA developed in a similar timeframe in both sexes. This is in agreement with previously published work, which found no sex differences in ethanol- or LiCl-induced CTA in adult male and female Long-Evans rats (Glover et al., 2016) or adult male and female Sprague Dawley rats (Vetter-O’Hagen et al., 2009). However, other work suggests that adult males develop stronger ethanol-induced CTA than adult females (Cailhol & Mormède, 2002; Morales et al., 2014; Schramm-Sapyta et al., 2014). Different methods for reporting saccharin intake could contribute to these discrepant findings. For example, though it may be tempting to normalize saccharin intake to body weight, previous work has shown that while males drink more than females overall, fluid intake is not directly related to body weight (T. Holdstock, 1973; Richter & Brailey, 1929; Santollo & Edwards, 2021). Therefore, absolute volume (mL) is likely a more accurate measure of intake than volume normalized to body weight (g/kg). Differential representation of rats that are sensitive or resistant to ethanol-induced CTA could also contribute to discordant results. Indeed, our previous work revealed sex differences in CTA only after rats were divided into distinct phenotypes, with differences being driven primarily by CTA-resistant females (Przybysz et al., 2023). Thus, it is possible that greater representation of a given phenotype in one sex over the other could contribute to the presence or absence of significant sex differences. Importantly, while a number of studies have considered sex differences in CTA acquisition and expression, no studies to date have examined incubation of CTA in female rats. Our data reveal that this phenomenon is observed to a similar degree in females as has been observed in males.

Findings from the present study suggest that previously published studies using CTA to measure tolerance to the aversive properties of ethanol were likely capturing the well-characterized US pre-exposure effect that can occur during Pavlovian conditioning. By eliminating the possibility of US pre-exposure using a within-subjects design, our data show that healthy augmentation of conditioned aversion is dampened after CIE exposure in both sexes (**Figure 7**). These results support existing data suggesting that neural circuits governing aversive signaling are disrupted after chronic ethanol exposure (Glover et al., 2019; Kang et al., 2017; J. Li et al., 2017; W. Li et al., 2023; Roberto et al., 2004, 2012). Such disruptions could contribute to maladaptive responding characteristic of AUD. The within-subjects design used in the present study can be used in future work to explore the neural mechanisms underlying this phenomenon.

**Figure 7.**
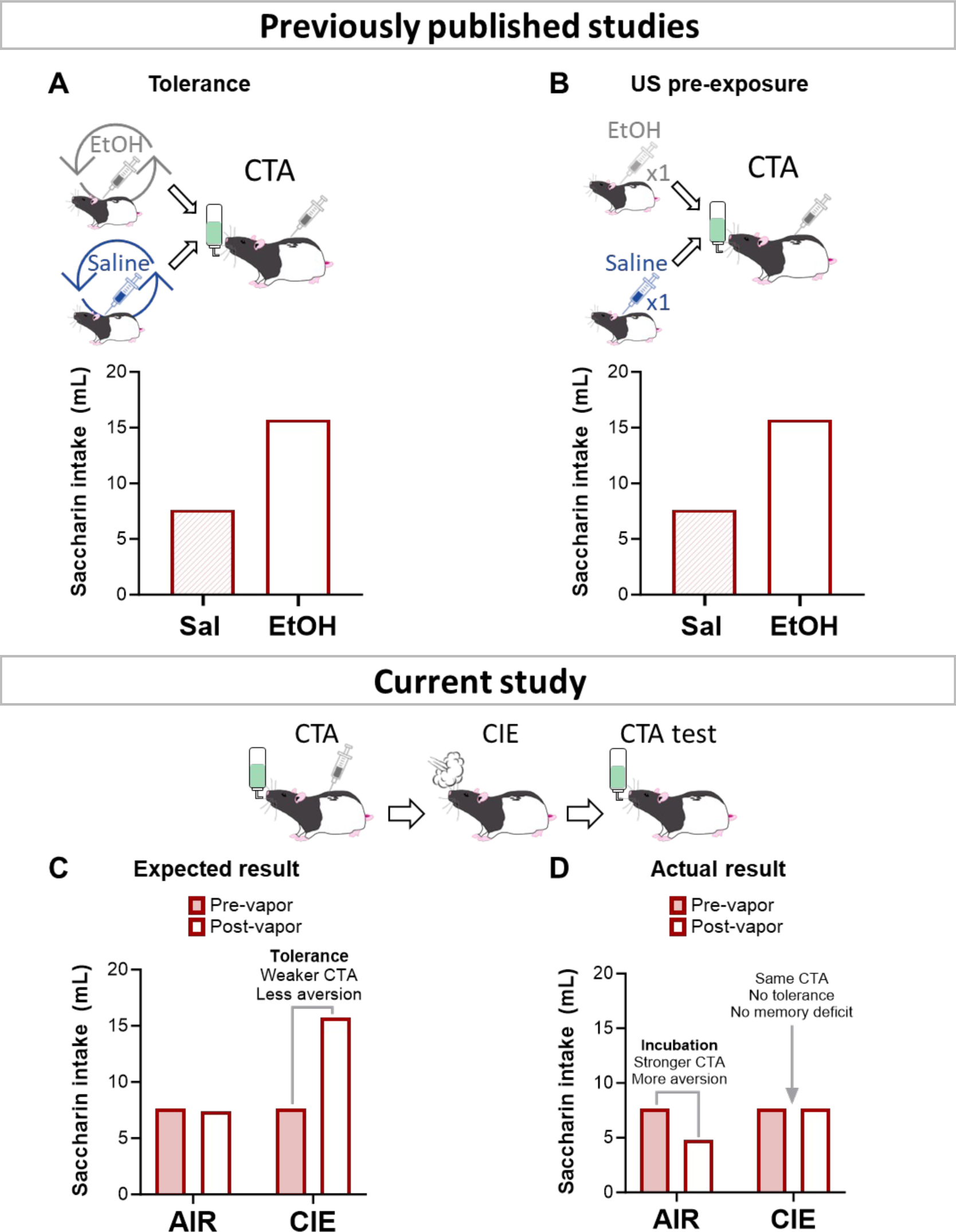
Within-subjects design uncovers attenuated incubation of EtOH-induced CTA after CIE exposure. Previously published work measured CTA following chronic ethanol exposure. Using this design, attenuation of CTA due to tolerance **(A)** is indistinguishable from attenuation of CTA due to US pre-exposure **(B)**. Using a within-subjects design that measures CTA before and after chronic ethanol exposure revealed unexpected findings. Instead of observing weaker CTA in CIE-exposed rats compared to pre-vapor exposure, indicative of tolerance **(C),** we found that CTA magnitude increased in AIR-exposed, but not CIE-exposed rats, indicative of attenuated incubation **(D)**.

## Supporting information

Supplemental results

## Acknowledgements

The authors would like to thank the following lab members for technical assistance: Christen Amegashie, Alex Brown, Nidhi Chetan, Kacey Clayton-Stiglbauer, Karl Bosque-Cordero, Andres Gascon, Shikun Hou, Nikki Kinarasri, Autumn Ollice, Arleen Perez Ayala, Jacqueline Sanchez, Shree Srinivasan, and Shannon Wheeler. This work was supported by NIH grants R01 AA029130 (EJG), P50 AA022538 (EJG), R00 AA024208 (EJG), T32 AA026577 (LAR; KRP). The authors have no financial or non-financial interests to disclose.

