## Supplemental results for "Attenuated incubation of ethanol-induced conditioned taste aversion in a model of dependence"

### Supplemental material

*The effect of vapor exposure on CTA is not different between phenotypes*

Our previous work uncovered the presence of two distinct CTA phenotypes, with some rats exhibiting increased sensitivity and others increased resistance to ethanol-induced CTA (Przybysz et al., 2023). To determine whether vapor exposure affected CTA magnitude differently in rats of each phenotype multifactorial repeated measures ANOVAs were performed comparing saccharin intake between phenotype and vapor group prior to and following vapor exposure. This analysis failed to uncover any effect of phenotype on the interaction between vapor exposure and time. Specifically, in males that received saccharin paired with saline, this analysis revealed a main effect of time [$F(1,22)=7.79$, $p=0.01$] and a time by vapor interaction [$F(1,22)=34.96$, $p<0.001$], but no three way interaction or two way interaction between vapor treatment and phenotype. Bonferroni-corrected multiple comparisons showed that, similar to results shown in **Figure 3**, CIE-exposed rats drank significantly more saccharin than AIR-exposed rats following vapor exposure ($p=0.006$, **Figure S1A**). For males injected with 1.5 g/kg EtOH, a main effect of time [$F(1,22)=5.18$, $p=0.03$] and of phenotype [$F(1,22)=33.42$, $p<0.001$] were observed but no three or two way interactions. Bonferroni-corrected multiple comparisons found that pre-vapor saccharin intake was significantly greater than post-vapor intake ($p=0.03$). In addition, as expected, rats in the resistant phenotype drank significantly more saccharin than rats in the sensitive phenotype ($p<0.001$, **Figure S1B**). For males injected with 2.0 g/kg ethanol, a significant three way interaction [$F(1,22)=5.18$, $p=0.03$] was observed as well as a significant interaction between time and phenotype [$F(1,22)=8.41$, $p=0.008$] and main effects of phenotype [$F(1,22)=53.93$, $p<0.001$] and of time [$F(1,22)=6.05$, $p=0.02$]. Because no main effect of vapor was observed, data were collapsed across vapor exposure and the interaction between time and phenotype was reassessed with a two-way ANOVA. This analysis revealed a significant interaction between time and phenotype [$F(1,24)=9.17$, $p=0.006$]. Sidak-corrected multiple comparisons found that rats in the resistant phenotype drank significantly less saccharin after vapor exposure than rats in the sensitive phenotype (p=0.0009, **Figure S1C**) regardless of vapor group.

The same analyses in females showed a significant main effect of phenotype [*F*(1,21)=9.08, p=0.007, **Figure S1D**] in the absence of any significant three or two way interactions. in rats that received saccharin paired with saline with rats in the sensitive phenotype consuming significantly more saccharin than rats in the resistant phenotype. Similarly, only a significant main effect of phenotype [*F*(1,20)=29.25, p<0.001, **Figure S1E**] was observed in females that received saccharin paired with 1.5 g/kg EtOH, with rats in the resistant phenotype drinking significantly more saccharin than those in the sensitive phenotype. While females that received saccharin paired with 2.0 g/kg EtOH did not have a significant three way interaction, there was a significant main effect of phenotype [*F*(1,21)=11.0, p=0.003] and a significant interaction between phenotype and time [*F*(1,21)=6.61, p=0.02]. In the absence of a main effect of vapor, data were collapsed across vapor group and a two-way ANOVA was run to assess the relationship between time and phenotype. This analysis showed a significant interaction between time and phenotype [*F*(1, 23)=15.14, p=0.0007]. Sidak-corrected multiple comparisons revealed that, similar to males that received saccharin paired with 2.0 g/kg EtOH, females in the resistant phenotype drank significantly less saccharin following vapor exposure compared to rats in the sensitive phenotype (**p=0.001, Figure S1F**). Altogether, these analyses show that while effects are less easily detected in sensitive rats due to a floor effect, CTA is similarly affected by vapor treatment in rats of both phenotypes.

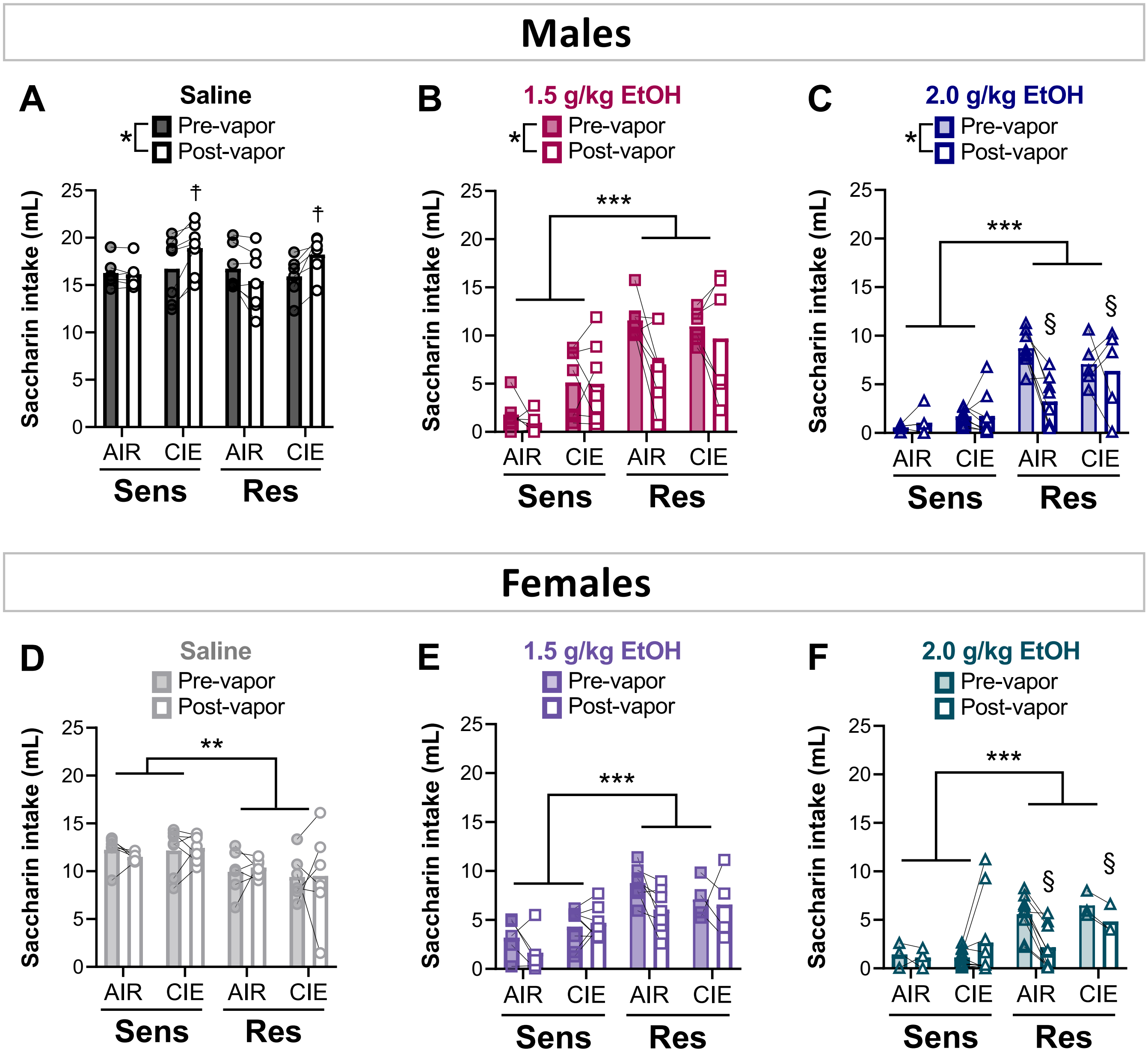


**Supplementary Figure 1. CTA is affected similarly by vapor in rats of both phenotypes.** There were no significant interactions between phenotype and vapor group in males **(A-C)** or females **(D-F)**. Males n=5-8 rats/group, females n=3-10 rats/group; $^{**}p<0.01$, $^{***}p<0.001$, $^{\dagger}p<0.01$ pre- vs post-vapor across vapor groups; $^{\S}p<0.001$ pre- vs post-vapor across phenotypes.
